# Expanding the palette of trehalose-based fluorophores for live mycobacterial detection

**DOI:** 10.64898/2026.07.16.737568

**Authors:** Adriann Brodeth, Reena Dosanjh, Lilith A. Schwartz, Sasha Kramer, Sean Goggins, Joel Cresser Brown, Abdulai Zigli, Benjamin M. Swarts, Mireille Kamariza

## Abstract

Tuberculosis (TB) remains the world’s leading infectious cause of death. Trehalose-based fluorogenic probes have emerged as powerful tools for labeling and studying mycobacteria, including *Mycobacterium tuberculosis* (Mtb), the causative agent of tuberculosis. However, existing probes occupy a limited spectral range and require compromise between brightness, specificity, and functional readouts. Here, we report the design and characterization of two trehalose conjugates derived from Janelia Fluor® dyes, JF_635_-Tre and JF_646_-Tre, which extend the trehalose-based platform into the far-red region. Following NHS ester-mediated synthesis, the trehalose-conjugated analogs displayed strong far-red fluorescence, with excitation/emission maxima at 638/654 nm and 648/663 nm for JF_635_-Tre and JF_646_-Tre, respectively. Both probes exhibited concentration- and time-dependent labeling of *Mycobacterium smegmatis* (Msmeg) and Mtb with minimal background fluorescence from the corresponding unconjugated dyes. Furthermore, we observed reduced labeling in heat-killed cells compared to live Msmeg, particularly for JF_646_-Tre, consistent with sensitivity to metabolic activity. Both JF-Tre derivatives produced significant cellular labeling and JF_635_-Tre distinguished untreated from INH-treated samples in drug susceptible Mtb, demonstrating the potential of the JF-Tre probes to report on INH susceptibility and resistance. Together, these findings expand the toolkit of trehalose-based probes and highlight how fluorophore identity influences probe performance in mycobacterial fluorescence imaging and drug susceptibility testing.

## Introduction

Tuberculosis (TB), caused by *Mycobacterium tuberculosis* (Mtb), is the leading infectious disease globally, with an estimated 10.8 million new infections and 1.25 million deaths reported in 2023.^1,2^ The growing prevalence of drug-resistant TB remains a major public health concern, with over 400,000 cases of rifampicin-resistant (RR) or multidrug-resistant (MDR) TB estimated in the same year.^1^ A key contributor to the persistence and drug tolerance of Mtb is its uniquely complex and lipid-rich cell envelope, which plays important roles in virulence, immune evasion, and intrinsic antibiotic tolerance.^3,4^ The outermost layer of this envelope, the mycomembrane, is composed largely of long-chain mycolic acids linked to trehalose, namely trehalose monomycolate (TMM) and trehalose dimycolate (TDM).^5–7^ These glycolipids are central to cell envelope assembly, host–pathogen interactions, and mycobacterial pathogenicity.^8–10^ Consequently, understanding the structure, dynamics, and biosynthesis of the mycomembrane is critical for both fundamental microbiology and the development of new diagnostic and therapeutic strategies.

Fluorescent chemical probes have emerged as valuable tools for visualizing bacterial growth and cell wall synthesis in living cells.^11^ Fluorescent D-amino acid derivatives have been widely used to label sites of peptidoglycan biosynthesis and monitor bacterial growth dynamics.^12–14^ Enzyme-activated probes have also been developed to report on specific enzymatic activities within the mycobacterial cell envelope, including probes targeting β-lactamase activity^15–17^, iron scavenging pathways^18^, protease activity^19^, or other cell-envelope–associated enzymes. While these approaches have enabled important insights into bacterial physiology, they primarily report on peptidoglycan or enzymatic activity rather than specifically labeling the mycomembrane. An alternative strategy exploits the distinctive role of trehalose within the mycomembrane. The antigen 85 (Ag85) protein complex catalyzes the transfer of mycolic acids onto trehalose substrates, a process that can be leveraged to metabolically incorporate trehalose analogs bearing various functional handles directly into the mycomembrane (**Figure 1A-1B**).^20,21^ Fluorine-^22,23^, azide-^24,25^, alkyne-^26,27^, fluorophore-functionalized^22,28–30^ trehalose moieties demonstrated successful metabolic incorporation into the mycomembrane and subsequent detection with fluorescence microscopy (**Figure 1B-1C**). However, many of these probes require multi-step labeling procedures, washing steps, or secondary reactions for signal generation. We and others have demonstrated that environment-sensitive dyes, such as 4-*N,N*-dimethylamino-1,8-naphthalimide trehalose (DMN-Tre)^31^, 3-hydroxychrome trehalose (3HC-Tre)^32^, and the far-red molecular rotor trehalose (RMR-Tre)^33^, have advanced the field by enabling wash-free imaging (**Figure 1C**). Similarly, FRET probes such as quencher-trehalose-fluorophore (QTF)^30^, FRET trehalose dimycolate (FRET-TDM)^34^, and arabinogalactan-linked mycolate (AGM)^35^ have shown wash-free labeling of mycobacteria. However, many current probes occupy a narrow spectral window, or exhibit trade-offs between brightness and labeling efficiency, restricting their utility in multiplexed assays designed to monitor different physiological processes simultaneously.

**Figure 1.**
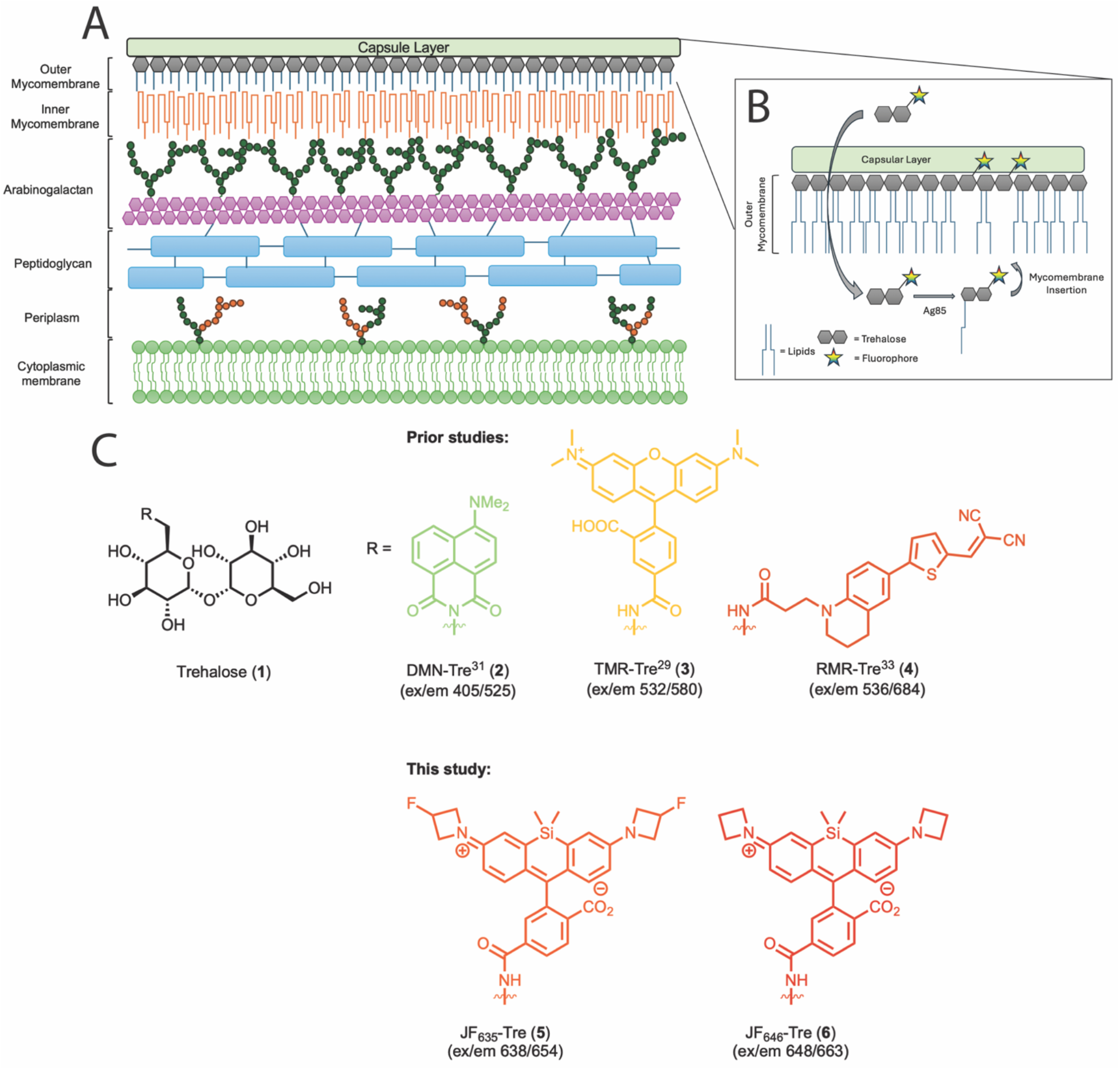
(**A**) Schematic representation of the cell envelope found in mycobacterial species. (**B**) Ag85-mediated metabolic incorporation of trehalose-based fluorophores into mycomembrane enabling rapid, specific mycobacterial labeling (adapted with permission from Kamariza, *et. al. Sci. Trans. Med*. eaam6310, 2018). (**C**) Structure of trehalose and trehalose probes used in this study, including JF_635_-Tre and JF_646_-Tre synthesized and characterized in this work. Colors correspond to emission wavelength.

To address these limitations, we sought to expand the mycobacterial chemical toolkit by incorporating high-performance, photostable fluorophores that operate in the far-red optical window. Janelia Fluor® (JF) dyes, derived from rhodamine, are renowned for their exceptional brightness, photostability, and tunable spectral properties that extend into the far-red region.^36–38^ Their compatibility with standard red-laser excitation makes them ideal for live-cell imaging, as it minimizes autofluorescence interference. In this work, we report the design and characterization of two novel trehalose conjugates derived from JF_635_ (JF_635_-Tre) and JF_646_ (JF_646_-Tre). We demonstrate that these probes are robustly incorporated into the mycomembrane of both *Mycobacterium smegmatis* (Msmeg) and Mtb in a concentration- and time-dependent manner. Additionally, we compare the labeling ability of JF_635_-Tre and JF_646_-Tre compared to other well-characterized trehalose probes (**Table 1**). We demonstrate robust labeling in different commercially available fluorescence filters by flow cytometry. Finally, we show that JF_635_-Tre probe reports on INH susceptibility and resistance. Together, these probes represent a significant expansion of the mycobacterial chemical toolkit into the far-red region, offering a sensitive platform for studying mycomembrane dynamics and accelerating the development of rapid diagnostic assays.

**Table 1.** Spectroscopic data and characterization of trehalose probes in this study categorized by emission wavelength.

| Fluorophore | Dye | MW | $\epsilon$ (c<sup>-1</sup>M<sup>-1</sup>) | Peak Excitation (nm) | Peak Emission (nm) |
| --- | --- | --- | --- | --- | --- |
| **Blue-to-Green** |  |  |  |  |  |
| 4-*N,N*-dimethylamino-1,8-naphthalimide | DMN-Tre<sup>31</sup> | 564.54 | 8,800 | 405 | 525 |
| **Green-to-Yellow** |  |  |  |  |  |
| TAMRA | TMR-Tre<sup>29</sup> | 753.75 | 90,000 | 532 | 580 |
| **Red** |  |  |  |  |  |
| Red molecular rotor turn-on | RMR-Tre<sup>33</sup> | 686.73 | 33,500 | 536 | 684 |
| Janelia Fluor® 635 | **JF<sub>635</sub>-Tre (this study)** | 855.92 | 130,000 | 638 | 654 |
| Janelia Fluor® 646 | **JF<sub>646</sub>-Tre (this study)** | 819.94 | 89,750 | 648 | 663 |

## Results and Discussion

To expand the spectral range of trehalose-based probes for mycobacterial labeling, we designed two trehalose conjugates derived from the JF dyes, JF_635_ and JF_646_ (**Figure 2**). We synthesized these JF-trehalose derivatives by coupling an amine-functionalized trehalose scaffold, 6-trehalosamine **1**, with commercially available NHS ester derivatives of JF_635_ and JF_646_, JF_635_ NHS ester **7** and JF_646_ NHS ester **8**, respectively, using standard amide coupling conditions (**Figure 2A**). Following purification by reverse-phase chromatography, the resulting probes, JF_635_-Tre **5** and JF_646_-Tre **6**, were obtained in good yield and characterized by NMR spectroscopy and high-resolution mass spectrometry.

**Figure 2.**
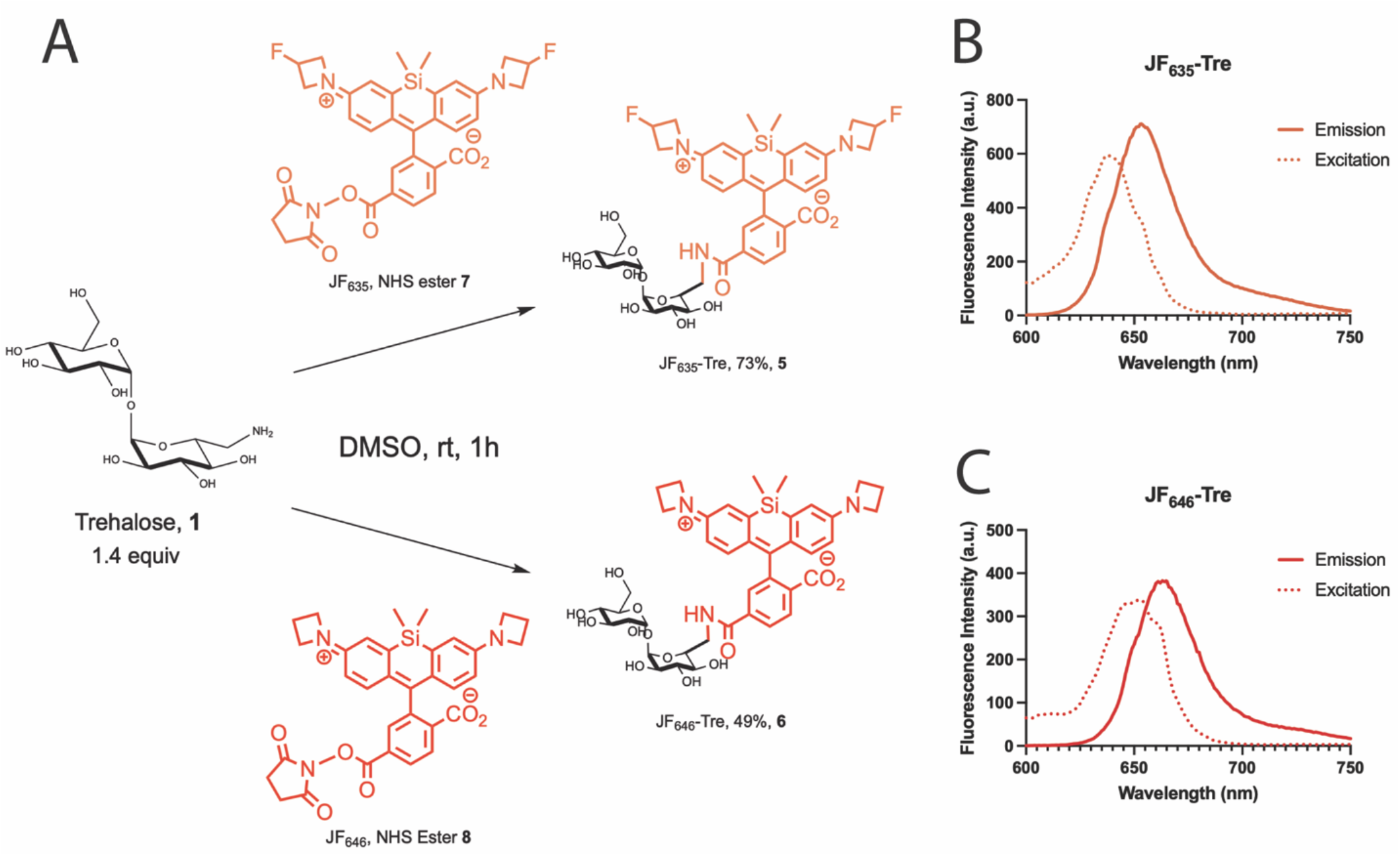
Synthesis scheme and fluorescence spectra for JF_635_-Tre and JF_646_-Tre. (A) Synthesis of JF_635_-Tre and JF_646_-Tre. Fluorescence emission and excitation spectra of (B) JF_635_-Tre and (C) JF_646_-Tre measured in 0.1% v/v trifluoroacetic acid (TFA) in 2,2,2-trifluoroethanol (TFE).

Using a fluorescence spectrophotometer, we measured and confirmed that both probes retain similar optical properties of the parent JF dyes. Indeed, JF_635_-Tre exhibited excitation and emission maxima at 638 and 654 nm, respectively, while JF_646_-Tre displayed excitation and emission maxima at 648 and 663 nm (**Figure 2B-C**). Both probes showed strong fluorescence in 0.1% v/v trifluoroacetic acid (TFA) in 2,2,2-trifluoroethanol (TFE) with spectral profiles consistent with previously reported JF derivatives.^36,37^ These properties position JF_635_-Tre and JF_646_-Tre within the far-red spectral window and make them suitable for fluorescence imaging using standard red-laser excitation.

Having established the probe properties, we sought to characterize the labeling properties of both JF_635_-Tre and JF_646_-Tre (**Figure 3**). We first evaluated whether JF_635_-Tre and JF_646_-Tre efficiently label mycobacteria. We treated Msmeg cultures with 10 µM JF_635_-Tre or 10 µM JF_646_-Tre, as well as the corresponding unconjugated, free dyes (JF_635_ Free, JF_646_ Free) for 4 hours at 37 ºC. Cells were then washed and analyzed by flow cytometry (**Figure 3A, 3H**). Both JF_635_-Tre and JF_646_-Tre produced significantly higher mean fluorescence intensity (MFI) compared to their unconjugated counterparts, which exhibited signal comparable to background (no dye and vehicle controls). Specifically, JF_635_-Tre and JF_646_-Tre achieved a ∼5-fold and ∼25-fold increase in signal, respectively.

**Figure 3.**
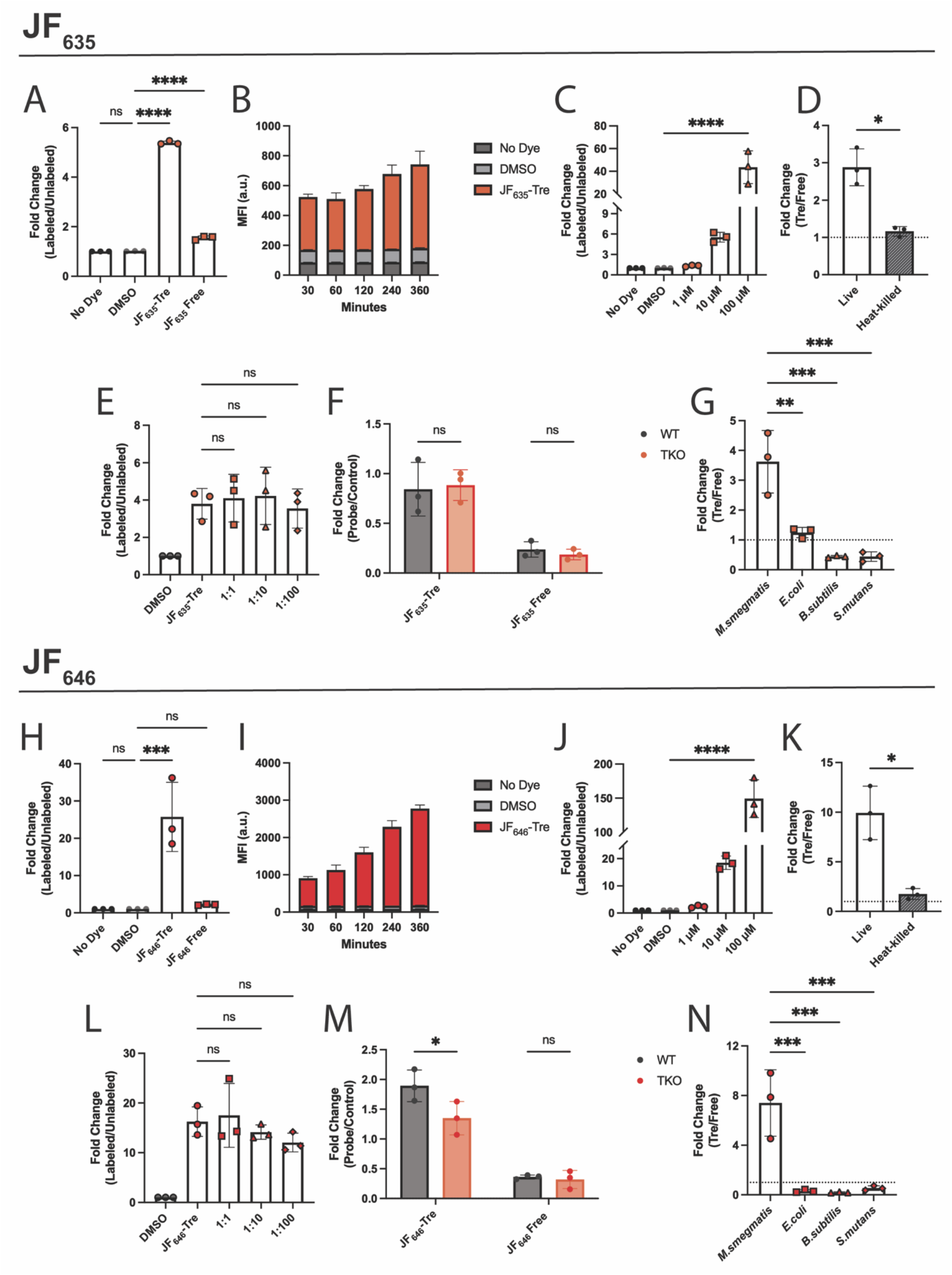
Characterization of JF_635_-Tre and JF_646_-Tre labeling of *Mycobacterium smegmatis* (Msmeg). JF_635_ (Upper Half): (**A**) Msmeg samples were incubated for 4 hours at 37 ºC with either 10 µM trehalose-conjugated JF_635_, 10 µM JF_635_ Free, 0.2% DMSO, or left untreated (No Dye). (**B**) Flow cytometry MFI analysis of Msmeg cells incubated with 10 µM JF_635_-Tre, 0.2% DMSO or left untreated (No Dye) for 30, 60, 120 or 360 minutes. (**C**) Fold-change analysis of flow cytometry MFI of Msmeg cells incubated with 1 µM, 10 µM, or 100 µM of JF_635_-Tre for 4 hours at 37°C. (**D**) Flow cytometry MFI analysis of live or heat-killed Msmeg cells incubated with 10 µM JF_635_-Tre or 10 µM JF_635_ Free for 1 hour at 37°C. (**E**) Flow cytometry MFI analysis of Msmeg cells incubated 1 hour with either 10 µM JF_635_-Tre, 10 µM JF_646_-Tre, or 0.2% DMSO for 1 hour at 37°C with increasing amounts of exogenous trehalose. (**F**) Flow cytometry MFI analysis of Msmeg cells incubated with either 10 µM JF_635_-Tre, 10 µM JF_646_-Tre or 10 µM RADA for 30 minutes at 37°C. (**G**) Labeling based on doubling times of Msmeg and non-mycobacterial species treated as in (**A**). **JF**_**646**_ (Lower Half): (**H-N**) Similar experiments were performed following the order in (**A-G**) replacing JF_635_ with JF_646_. MFI (a.u.) = Mean Fluorescence Intensity (arbitrary units). Error bars denote standard deviation of three biological replicates and data were analyzed by t-test or ANOVA tests in GraphPad Prism. *p* values: * < 0.05, ** < 0.01, *** < 0.001, **** < 0.0001.

Next, we aimed to characterize the Msmeg labeling parameters. We performed a time-course analysis by incubating Msmeg cultures with 10 µM JF_635_-Tre or 10 µM JF_646_-Tre at 30, 60, 120, 240, or 360 minutes at 37 ºC before flow cytometry analysis (**Figure 3B, 3I**). Both JF_635_-Tre and JF_646_-Tre exhibit progressive increases in fluorescence intensity over the course of incubation. Consistent with our previous results, JF_646_-Tre produced a more pronounced fluorescence increase compared to JF_635_-Tre. Both probes produced clear signals above background within 30 minutes, demonstrating rapid trehalose-mediated uptake. Similarly, we observed that JF-Tre labeling also increased in a dose-dependent manner (1-100 µM), with significant signal observed at concentrations as low as 10 µM (**Figure 3C, 3J**). JF_646_-Tre consistently produced higher fluorescence values across all concentrations tested, whereas JF_635_-Tre showed lower overall signal but maintained clear concentration-dependent behavior. Overall, the JF-Tre probes, especially JF_646_-Tre, achieve strong labeling with short incubation times and modest probe quantities, enabling rapid mycobacterial detection. In addition, their red-shifted excitation (635 and 646 nm) and emission (670 and 710 nm) wavelengths reduced background autofluorescence and yielded robust signal-to-noise ratios.

To test whether metabolic activity is required for labeling, we subjected live or heat-killed Msmeg to JF-Tre probe labeling. We incubated Msmeg cells under normal growth conditions and then subjected to heat-killing (95 ºC for 30 minutes) followed by free and trehalose-conjugated probe labeling as before (**Figure 3D, 3K, Figure S1**). We observed strong labeling of live Msmeg with JF_646_-Tre compared to heat-killed Msmeg, whereas JF_635_-Tre only had a modest fluorescence difference between live and heat-killed Msmeg (**Figure S1**). While we observed a modest reduction for JF_635_-Tre, we observed similar fluorescence signals when incubated with the unconjugated free dye, suggesting likely nonspecific dye interaction leading to turn on. By taking the ratio of fluorescence signal from the trehalose-conjugated form over its corresponding unconjugated acid, untreated samples show ∼3-fold and ∼10-fold increase for JF_635_-Tre and JF_646_-Tre, respectively (**Figure 3D, 3K**). In contrast, heat-killed bacteria demonstrated a fluorescence ratio of ∼1 for both JF probes indicating that labeling due to unconjugated JF dyes does not differentiate between live or heat-killed Msmeg. Ultimately, while some residual signal was observed, the overall reduction in JF_646_-Tre fluorescence indicates that this probe labeling is strongly associated with cellular viability and metabolic activity.

To characterize probe specificity, we tested whether JF-Tre incorporation depends on the trehalose metabolic pathway by labeling Msmeg in the presence of increasing amounts of excess exogenous trehalose (1:1, 1:10, or 1:100 of JF-Tre:trehalose) (**Figure 3E, 3L**). JF_646_-Tre fluorescence demonstrated limited non-significant decrease as trehalose concentrations increased (**Figure 3L**). Similarly, the JF_635_-Tre showed no substantial reduction at any ratio (**Figure 3E**). In addition, unconjugated forms of both dyes showed no change in fluorescence at any ratio (**Figure S2A-B**). By further analysis, at a 1:100 probe to trehalose ratio, JF_646_-Tre labeling decreased by nearly 25%, while JF_635_-Tre decreased by only 5%, indicating weaker sensitivity to trehalose competition compared to JF_646_-Tre and other previously published trehalose probes^31–33^ (**Figure S3A-D**). These results suggest limited dependence on trehalose-associated pathways for the two probes, suggesting that differences in fluorophore structure may play a role in the permeability and nonspecific turn-on behavior of the probes.

To determine whether JF-Tre probes are specifically incorporated via trehalose metabolic pathway, we assessed labeling in an antigen85 (Ag85) partial knockout mutant, *ΔMSMEG_6396-6399*, which lacks three of five genes in Msmeg putatively encoding Ag85 mycolyltransferases^31^. We used a control compound that labels peptidoglycan, a TAMRA-based fluorescent D-amino acid (RADA), to account for possible effects of Ag85 depletion on cell envelope permeability.^39^ By normalizing the fluorescence of the JF analogues to the RADA control, we observed a decrease in incorporation with JF_646_-Tre in the mutant bacteria (**Figure 3M**). In contrast, JF_635_-Tre and both unconjugated JF probes all show no significant difference between the wild-type and knockout mutant (**Figure 3F**). Although the data suggests JF-Tre probes are not primarily incorporated via trehalose metabolic pathways, these results highlight the potential of JF-Tre probes to distinguish live from dead mycobacteria.

Next, we evaluated the JF-Tre probe selectivity for mycobacteria against multiple bacterial species. We compared Msmeg, which possesses a mycomembrane, to Gram-negative *E. coli* and Gram-positive *B. subtilis* and *S. mutans*, which lack a mycomembrane, by treating each organism with 10 µM of either trehalose-conjugated probe or their free acid control compound and analyzing by flow cytometry (**Figure 3G, 3N, Figure S4**). As expected, both JF_635_-Tre and JF_646_-Tre efficiently labeled Msmeg **(Figure 3G, 3N**). While JF_646_-Tre demonstrated limited labeling of *E. coli, B. subtilis*, and *S. mutans*, all mycomembrane-deficient species, JF_635_-Tre displayed increased fluorescence signal with *E. coli*, and a mild fluorescence increase with *B. subtilis* (**Figure 3G, 3N**). Although JF_635_-Tre labeled both Msmeg and *E. coli* at similar levels, unconjugated JF showed low fluorescence in Msmeg but increased labeling in non-mycobacterial species (**Figure S3**). Thus, these results show that trehalose conjugation of the JF dyes enhanced their mycobacterial specificity.

We compared the JF probe labeling performance to previously reported trehalose probes, including DMN-Tre, TMR-Tre, and RMR-Tre, using flow cytometry across multiple fluorescence channels corresponding to standard excitation/emission settings (**Figure 4A-4B, Figure S5**).^31–33^ We incubated Msmeg and Mtb H37Ra in standard growth conditions to optical density (OD_600_) of 0.5, then incubated the cultures with 10 µM JF_635_-Tre, 10 µM JF_646_-Tre, 100 µM RMR-Tre, 100 µM DMN-Tre, and 100 µM TMR-Tre. Fluorescence values were obtained using 488 nm blue excitation laser for the 533/30 nm (FITC), 585/40 nm (PE), and 670 nm (PerCP) emission filters, optimized for DMN-Tre, TMR-Tre, and RMR-Tre, respectively. Additionally, fluorescence values were also obtained using 640 nm red excitation lasers for the 675/25 nm (APC) emission filter, optimized for the JF-Tre probes. For both strains, DMN-Tre, TMR-Tre and RMR-Tre produced strong fluorescence in their respective emission wavelength channels, consistent with their established spectral properties^33,40,41^ (**Figure 4A-B**). In contrast, JF_646_-Tre exhibited the highest fluorescence intensity in the APC channel, corresponding to far-red excitation (640 nm) and emission (675 nm), with minimal cross-talk in lower wavelength channels. JF_635_-Tre also produced signal in the far-red region, although at lower intensity compared to JF_646_-Tre. Importantly, signal from JF probes was largely restricted to the far-red channel, whereas other probes showed broader spectral distribution or peak intensity in shorter wavelength channels (**Figure 4A-B**). These results demonstrate that JF_635_-Tre and JF_646_-Tre extend the trehalose probe platform into the far-red spectral window, enabling compatibility with red-laser excitation and minimal overlap with probes occupying lower wavelength channels.

**Figure 4.**
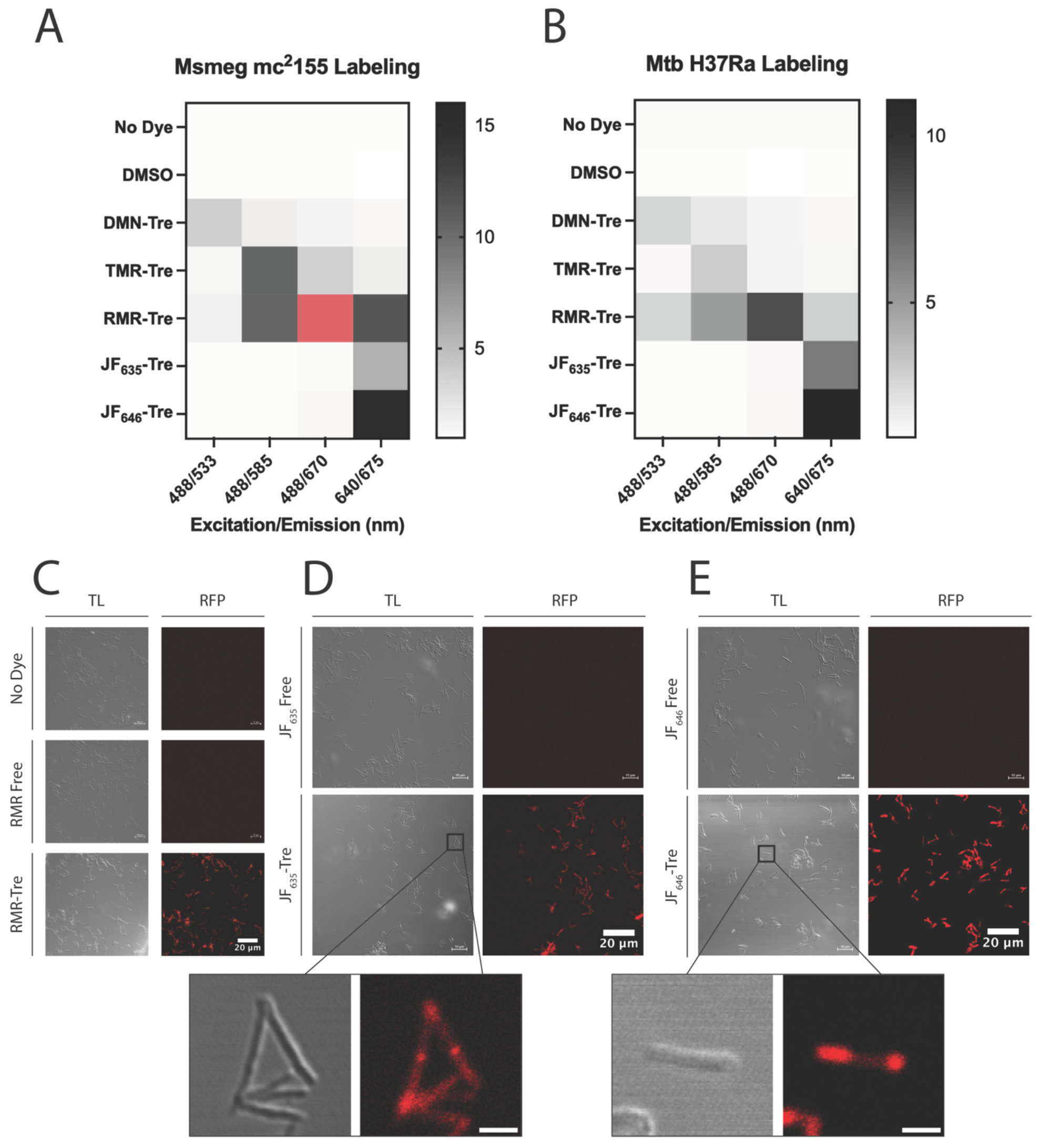
Assessment of JF-Tre probe labeling of Msmeg and *Mycobacterium tuberculosis* (Mtb) compared to existing trehalose probes. (**A-B**) Heatmap representations of fluorescence fold-change over no dye controls. (**A**) Msmeg cultures were incubated for 4 hours and (**B**) Mtb cultures were incubated for 16 hours at 37 ºC with 100 µM DMN-Tre, 100 µM TMR-Tre, 100 µM RMR-Tre, 10 µM JF_635_-Tre, 10 µM JF_646_-Tre, 0.2% DMSO, or left untreated (No Dye). Fluorescence values were obtained using 488 nm blue excitation laser for the FITC (533/30 nm), PE (585/40 nm), and PerCP (670 nm) emission filters and 640 nm red excitation lasers for the APC (675/25 nm) emission filter. Red box in Figure **4A** represents a mean fold change = 108.06. Values are means from three biological replicates. (**C-E**) Representative confocal microscopy images of Msmeg cells taken after a 4-hour incubation at 37 ºC in the presence of (**C**) RMR Free, RMR-Tre, (**D**) JF_635_ Free, JF_635_-Tre, (**E**) JF_646_ Free, or JF_646_-Tre. Cells were washed in DPBS and imaged using a Nikon A1R Laser Scanning Confocal Microscopy. MFI (a.u.), mean fluorescence intensity (arbitrary units). Scale bar = 2.5 µm, unless noted otherwise. TL = transmitted light.

To validate probe labeling at the cellular level, Msmeg cells were imaged by fluorescence microscopy following incubation with representative probes. Cells treated with RMR-Tre exhibited strong fluorescence localized to the cell periphery, consistent with labeling of the mycobacterial cell envelope, whereas untreated controls and cells treated with free RMR dye showed minimal fluorescence (**Figure 4C**). Similarly, JF_635_-Tre and JF_646_-Tre produced clear fluorescence signals associated with bacterial cells, while the corresponding unconjugated dyes (JF_635_ Free and JF_646_ Free) resulted in negligible signal (**Figures 4D-E**). In both cases, fluorescence appeared localized to the polar regions of the cell, consistent with the polar growth of mycobacteria and labeling of the cell envelope. Notably, JF_646_-Tre produced a more intense fluorescence signal compared to JF_635_-Tre, also shown with flow cytometry quantitative measurements in **Figure 3A** and **3H**. Nonetheless, both probes enabled visualization of individual bacterial cells with minimal background from free acid dye controls.

Due to their established dependence on metabolic activity, DMN-Tre and RMR-Tre have previously been shown to report on drug susceptibility, enabling detection of Mtb strain responses to drug treatment.^31,33,42^ Thus, we assessed the potential of JF-Tre probes to monitor Mtb drug susceptibility using isoniazid (INH), a first-line anti-TB drug that inhibits mycolic acid biosynthesis. Here, we tested INH-resistant strains of Msmeg (*ΔkatG*) and Mtb H37Rv (mc^2^8245), along with their drug-susceptible counterparts wild-type Msmeg mc^2^155 and Mtb H37Rv (mc^2^7902) (**Figure 5**). Msmeg cultures were first treated with 10 µg/mL INH for 24 hours then labeled with 10 µM JF-Tre. We found that both JF_635_-Tre and JF_646_-Tre distinguished untreated from INH-treated samples in drug-susceptible Msmeg, while INH-resistant strains showed no significant fluorescence change (**Figure 5A-C**). Similarly, Mtb cultures were treated with 1 µg/mL INH before adding JF-Tre probes and extended probe incubation to 16 hours to accommodate the slower metabolic rate of Mtb. We found that JF_635_-Tre distinguished untreated from INH-treated samples in drug susceptible Mtb, while JF_646_-Tre did not produce statistically significant fluorescence reduction (**Figure 5D-E**). Nonetheless, both probes showed no significant fluorescence change with INH-resistant strains. Overall, these findings demonstrate the potential of the JF-Tre probes to report on INH susceptibility and resistance and may be valuable for drug screening platforms due to their high signal-to-background ratio.

**Figure 5.**
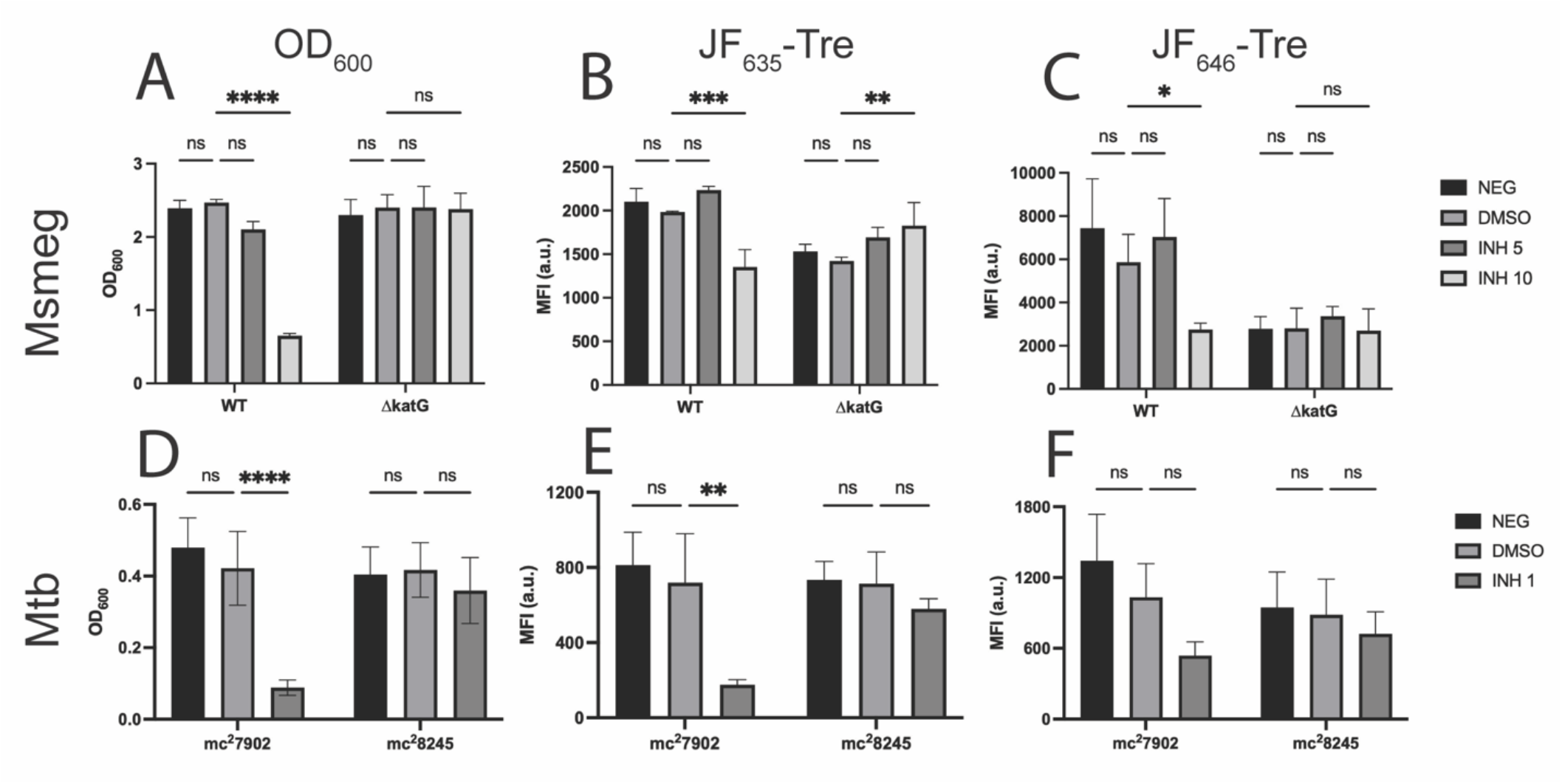
Isoniazid (INH) drug susceptibility in Msmeg and Mtb using JF_635_-Tre and JF_646_-Tre. (**A-C**) INH-susceptible (WT) and INH-resistant (*ΔkatG*) Msmeg strains were incubated for 20 hours treated with 5 or 10 µg/mL INH, 1% DMSO, or left untreated (NEG). Cultures were then incubated for an additional 4 hours with 10 µM JF_635_-Tre or 10 µM JF_646_-Tre, washed three times with PBS, and analyzed by fluorescence plate reader for (**A**) Optical density at 600 nm (OD_600_); and by (**B-C**) flow cytometry on the APC channel (640/675 nm, excitation/emission). (**D-F**) INH-susceptible (mc^2^7902) and INH-resistant (mc^2^8245) Mtb strains were incubated for 3 days at 37 ºC treated with 1% DMSO, 1 µg/mL INH, or left untreated (NEG). Cultures were incubated for additional 16 hours with either 10 µM JF_635_-Tre or 10 µM JF_646_-Tre, washed three times with PBS, fixed, and analyzed by plate reader and flow cytometry as in (**A-C**). MFI (a.u.) = Mean Fluorescence Intensity (arbitrary units). Error bars denote standard deviation of three biological replicates and data were analyzed by ANOVA tests in GraphPad Prism. *p* values: * < 0.05, ** < 0.01, *** < 0.001, **** < 0.0001.

## Conclusion

TB continues to impose a significant global burden, highlighting the urgent need for rapid, reliable diagnostic tools to limit TB transmission and enable timely treatment. Trehalose-based fluorophores represent a simple yet effective strategy for the specific and sensitive detection of mycobacteria, serving as powerful tools for both elucidating fundamental mycobacterial biology and developing low-cost diagnostics. However, existing trehalose probes occupy a broad, overlapping spectral window, or involve trade-offs between brightness and labeling efficiency, limiting the options available for diverse imaging and clinical applications. In this work, we expand this toolkit by introducing JF-conjugated probes, JF_635_-Tre and JF_646_-Tre, which extend trehalose-based detection into the deep-red excitation and emission range. Our results show that both probes generate strong signal-to-background ratios restricted in the far-red range, matching the performance of established conjugates while opening up new spectral territory. Importantly, their selective fluorescence turn-on in the far-red region provides the necessary photophysical foundation for potential multiplexed labeling experiments for mycobacteria. Indeed, our research demonstrates that JF_646_-Tre stands out as a highly robust tool, displaying exceptional brightness and specificity, while both derivatives demonstrate utility in various applications such as microscopy and drug-susceptibility testing. Future work will focus on deploying these probes in co-labeling systems alongside shorter-wavelength fluorophores, as well as exploring additional JF variants to span the visible spectrum more fully, enabling greater flexibility and control in fluorescence-based studies.

## Materials and Methods

### General Procedures for Compound Synthesis and Characterization

Reagents and solvents were procured from commercial sources without further purification unless otherwise noted. Purification by prep-HPLC was performed using an Agilent 1100 system with a Waters Atlantis T3 5μm 19 × 150mm column. Analytical thin layer chromatography was performed on glass plates pre-coated with silica gel Analtech, UNIPLATE− 250μm / UV254), with visualization being achieved using UV light (254 nm) and/or by staining with either alkaline potassium permanganate dip, acidic ninhydrin dip, or charring with 5% H_2_SO_4_ in EtOH. Nuclear magnetic resonance spectra were recorded on a Bruker Avance III HD spectrometer. High resolution mass spectral (HRMS) data was collected using an Agilent 6545 LC/Q-TOF system. Normalized absorption and fluorescence emission spectra were recorded in 10mM PBS pH 7.3 at the concentration noted for each sample following dilution of a DMSO stock solution. Absorption spectra were recorded with an Agilent Cary 60 UV-Vis spectrophotometer using genuine precision quartz cells from Lovibond with a 1 cm path length. Fluorescence spectra were recorded on an Agilent Cary Eclipse Fluorescence Spectrophotometer using either precision Quartz Suprasil cells from Hellma Analytics or Brand Disposable Cuvettes and a 1 cm path length.

### Bacterial Strains, Media, and Reagents

The bacterial strains used in this work included *M. smegmatis* mc^2^155, *M. smegmatis* ΔMSMEG_6396-6399^31^, *M. smegmatis* ΔkatG^42^, auxotrophic strains of *M. tuberculosis* H37Rv mc^2^7902 (drug-susceptible, ΔpanCD ΔleuCD ΔargB) and mc^2^8245 (INH-resistant, ΔpanCD ΔleuCD ΔargB), *Escherichia coli (E. coli), Bacillus subtilis (B. subtilis)*, and *Streptococcus mutans (S. mutans). M. smegmatis* and *M. tuberculosis* were cultured in Middlebrook 7H9 liquid medium supplemented with 10% (v/v) OADC, 0.5% (w/v) glycerol, and 0.05% (w/v) Tween 80. ΔMSMEG_6396-6399 culture medium was supplemented with 50 µg/mL hygromycin. *M. tuberculosis* culture medium was further supplemented with 24 mg/L pantothenate, 50 mg/L L-leucine, and 200 mg/L arginine. 5 µL of vitamins, dissolved in water, were added per 1 mL of 7H9 media. *E. coli* and *B. subtilis* were cultured in LB liquid medium, while *S. mutans* was cultured in Brain-Heart Infusion liquid medium. All bacteria were cultured at 37°C.

Stock solutions of DMN-Tre (20 mM) were prepared in water and stored at -20°C. Stock solutions of RMR-Tre and TMR-Tre were prepared in DMSO (20 mM) and stored at -20°C. Similarly, both the trehalose-conjugated and free acid forms of JF_635_ and JF_646_ were dissolved in DMSO (20 mM) and stored at -20°C. Before usage in labeling experiments, stock solutions were diluted to the desired concentrations.

### General growth and labeling procedure for bacteria

Bacterial cultures were started by inoculating from frozen bacterial stocks into 10 mL of liquid medium in a 50 mL culture tube. Starter cultures were incubated overnight at 37°C with shaking, reaching logarithmic phase. Before any experiments, cultures were diluted with liquid medium to the desired OD_600_ concentration. Labeling experiments were performed in aerated culture tubes. Probe stock solutions were inoculated into respective tubes. The final DMSO concentration in each solution (typically with 1.5 mL final volume) was 0.2%. The tubes were then incubated at 37°C with shaking until the desired endpoint. For wash steps, 1 mL of incubated and labeled culture was well suspended and transferred into a 1.5 mL Eppendorf tube. The tubes were centrifuged (21300 xg, 1 min, room temperature) and washed with 1x Dulbecco’s Phosphate Buffer Saline (DPBS) twice. An additional wash step (10000 xg, 5 min, room temperature) was performed and samples were resuspended in DPBS (for flow analysis and fluorescence plate reader analysis) or in deionized water (for microscopy).

### Flow cytometry analysis

After fluorescence labeling, 150 µL of sample was aliquoted into an opaque 96-well plate for flow and fluorescence plate reader analysis, done in quadruplicate for each biological replicate. Flow cytometry was performed on a BD Biosciences C6 Accuri™ Plus flow cytometer. This instrument is equipped with 488-nm blue and 640-nm red excitation lasers. This instrument also comes with 4 standard optical filters: FITC (533/30 nm), PE (585/40 nm), PerCP (>670 nm), and APC (675/25 nm). Fluorescence data were collected for 100,000 events per sample using a minimum threshold (10 FSC-H) for drug experiments and bacteria-only threshold (2000 FSC-H) for specificity experiments. Flow cytometry data was processed using BD Accuri™ C6 Plus Software and graphing and data analysis were done using GraphPad Prism.

### Concentration dependence of labeling

*M. smegmatis* cultures at an optical density at 600 nm (OD_600_) of 0.5 in 7H9 liquid medium were treated with increasing concentrations of either JF_635_-Tre or JF_646_-Tre. 0.2% DMSO and untreated cultures were also included as controls. Samples were incubated with shaking at 37°C for 4 hours. The cells were pelleted by centrifugation, washed, re-suspended in PBS, and analyzed by flow cytometry.

### Time dependence of labeling

*M. smegmatis* cultures at OD_600_ of 0.5 in 7H9 liquid medium were treated with either JF_635_-Tre or JF_646_-Tre at a final concentration of 10 µM. 0.2% DMSO and untreated cultures were also included as controls. Samples were incubated shaking at 37°C and taken out at the indicated time points. The cells were pelleted by centrifugation, washed, re-suspended in PBS, and analyzed by flow cytometry.

### Heat-killing

Two large aliquots of *M. smegmatis* cultures at OD_600_ of 0.5 in 7H9 liquid medium was prepared prior to heat-killing. One aliquot was heated at 95°C for 30 min to heat-kill bacteria, while the other was left unheated. Next, 1.5 mL of heat-killed and live bacteria were aliquoted into 5 mL culture tubes and treated with either JF-Tre or JF Free at a final concentration of 10 µM. 0.2% DMSO and untreated cultures were also included as controls. Samples were incubated with shaking at 37°C for 4 hours. The cells were pelleted by centrifugation, washed, re-suspended in PBS, and analyzed by flow cytometry.

### Trehalose competition

*M. smegmatis* cultures at OD_600_ of 0.5 in 7H9 liquid medium were treated with either JF-Tre or JF Free acid at a final concentration of 10 µM. 0.2% DMSO and untreated cultures were also included as controls. Afterwards, trehalose was added with final concentrations of 10, 100 or 1000 µM or left untreated (no trehalose). Samples were incubated shaking at 37°C for 1 hour. The cells were pelleted by centrifugation, washed, re-suspended in PBS, and analyzed by flow cytometry. JF_635_ and JF_646_ experiments were done on separate days.

### Labeling in Ag85 triple knockout mutant

*M. smegmatis*, mc^2^155 and ΔMSMEG_6396-6399, a triple knockout mutant of Ag85, at OD_600_ 0.5 in 7H9 liquid medium were treated with 0.05% DMSO (solvent control), 10 µM JF-Tre of JF Free acid (0.05% DMSO final concentration), 100 µM RMR-Tre or TMR-Tre (0.05% DMSO final concentration), 100 µM DMN-Tre, or 10 µM RADA (0.02% DMSO final concentration), and untreated cultures were also included as controls. Samples were incubated shaking at 37ºC for 30 minutes. The cells were washed three times in DPBS, as previously described, and analyzed by flow cytometry.

### Bacterial species specificity of labeling

*M. smegmatis* cultures at OD_600_ of 0.5 in 7H9 liquid medium, *E. coli* and *B. subtilis* cultures at OD_600_ of 0.5 in Luria Broth liquid medium, and *S. mutans* cultures at OD_600_ of 0.5 in Brain-Heart Infusion liquid medium were treated with 10 µM JF-Tre or JF Free acid. Untreated cultures and 0.5% DMSO treated cultures were also included as controls. *M. smegmatis* samples were incubated shaking at 37°C for 4 hours, *E. coli* and *B. subtilis* samples were incubated shaking at 37°C for 30 minutes, and *S. mutans* samples were incubated shaking at 37°C for 1.5 hours. Following incubation, the cells were pelleted by centrifugation, washed, re-suspended in DPBS, and analyzed by flow cytometry as previously described.

### Drug susceptibility assays

For *M. smegmatis*, drug-susceptible (mc^2^155) and drug-resistant (*Δ*katG) cultures were diluted to an OD_600_ of 0.1 in 5 mL aliquots. Cultures were inoculated with Isoniazid (INH) at a final concentration of 5 and 10 µg/mL. Controls included untreated and 0.1% DMSO. Cultures were incubated at 37°C for 20 hours, followed by incubation in 10 µM JF-Tre for an additional 4 hours. The cells were pelleted by centrifugation, washed for three times, and re-suspended in PBS, and analyzed by flow cytometry.

For *M. tuberculosis*, drug-susceptible (*mc*^*2*^*7902)* and drug-resistant (*mc*^*2*^*8245*) cultures were diluted to an OD_600_ of 0.1 in 4 mL aliquots. Cultures were inoculated with INH at a final concentration of 1 µg/mL. Controls included untreated and 1% DMSO. Cultures were incubated at 37°C for 3 days, followed by incubation in 10 µM JF-Tre for an additional 16 hours. The cells were pelleted by centrifugation, washed for three times, re-suspended in PBS, and analyzed by flow cytometry.

### Fluorescence microscopy

*M. smegmatis* cultures were generated by inoculating a single isolated colony from a freshly streaked LB agar plate into 3 mL liquid medium in a culture tube. Cultures were incubated at 37 °C with shaking until reaching mid-logarithmic phase, then diluted with liquid medium to an optical density at 600 nm (OD_600_) of 0.4 to initiate the experiment. 198 µL of the cell suspension were transferred into the wells of a 96-well plate and treated with 2 µL of RMR-Tre, RMR-Free, JF_635_-Tre, JF_635_-Free, JF_646_-Tre, and JF_646_-Free (all stocks 10 mM in DMSO), for a total volume of 200 µL per well. The final concentration of probes was 100 μM and the final concentration of DMSO was 1%. Plates were incubated at 37 °C with shaking in a Tecan plate reader (Infinite F200 PRO operated by Tecan iControl software) for 4 h. For wash steps, cells were transferred to a v-bottom 96-well plate, centrifuged (3,200 xg, 5 min, room temperature), and washed with PBS 1x containing 0.5% bovine serum albumin (PBSB) twice. The cells were then fixed in 4% formaldehyde in PBS and washed once with PBS. Finally, cells were prepared for analysis using flow cytometry or fluorescence microscopy.

For confocal microscopy analysis, 10 μL aliquots of labeled cells were placed directly onto a microscope slide, lightly spread into a thin layer using the edge of a coverslip, and briefly air-dried in the dark. Fluoromount-G mounting medium (SouthernBiotech) was applied, then coverslips were placed over the sample and immobilized with adhesive. Microscopy was immediately carried out using a Nikon A1R Laser Scanning Confocal Microscopy LSCM equipped with a 60X/1.40 NA Plan Apo oil-immersion objective lens with DIC optics. Images were acquired using two lasers and two detectors. RMR-Tre was excited using the 561 nm diode laser, and fluorescence emission was collected using the Alexa Fluor detector channel (em = 684 nm). JF_635_-Tre and JF_646_-Tre were excited using the 640 nm diode laser, and fluorescence emission was collected using the Cy5.5 detector channel, appropriate for their respective excitation and emission wavelengths (JF_635_-Tre: em = 675 nm; JF_646_-Tre: em = 710 nm). Image acquisition and processing were performed using Nikon NIS-Elements AR, with the same post-processing applied to all images using FIJI.

## Acknowledgements

We thank the Jacobs Lab at Albert Einstein College of Medicine and the Bertozzi Lab at Stanford University for their generous donations of auxotrophic Mtb strains, and Msmeg *ΔkatG*, respectively. This work was supported by a UC LEADS scholarship to A.B and a NIH NIGMS T32 GM136614 predoctoral fellowship to L.A.S. This research was supported by a NIH R01 AI179891 grant to M.K.

## Author contributions

J.C.B., B.M.S., and M.K. designed and led the study. A.B., R.D., L.A.S., S.K., S. G., and A. Z. performed most of the experiments. A.B. and M.K. analyzed all of the data. A.B. and M.K. wrote and edited the manuscript, which was approved by all authors.

