## Supplemental Material for "Expanding the palette of trehalose-based fluorophores for live mycobacterial detection"

**The PDF file includes**

**Materials and Methods**
- Synthesis of $JF_{635}$-Tre
- Synthesis of $JF_{646}$-Tre

**Figures**
- **Fig. S1**. Raw Fluorescence Data of Live vs. Heat-killed Labeling
- **Fig. S2.** Trehalose Competition for Unconjugated $JF_{635}$ and $JF_{646}$
- **Fig. S3.** Percentage Change Analysis for JF-Tre Trehalose Competition
- **Fig. S4.** Raw Fluorescence Data of Bacterial Specificity Labeling
- **Fig. S5.** Fluorescence Analysis of Msmeg and Mtb labeling with different trehalose probes

**Appendix**
- NMR spectra of $JF_{635}$-Tre
- NMR spectra of $JF_{646}$-Tre

### Materials and Methods

**Synthesis of** $JF_{635}$**-Tre**

To a stirring solution of 6-trehalosamine (38mg, 0.11mmol, 1.4 equiv.)[43] in DMSO (4mL) was added $JF_{635}$ NHS ester (50mg, 79μmol, 1 equiv.).[37] The reaction was stirred at room temperature for 1 hour before being purified directly by prep-HPLC (10–90% MeCN/0.1% formic $acid_{(aq.)}$). Pure product fractions were then combined and lyophilized to give the title compound as a light blue solid (50mg, 73%).

$^{1}$H NMR (400 MHz, $d_4$-MeOD); $\delta_H$ 8.04 (1H, dd, $J$ = 8.0, 1.3 Hz), 8.01 (1H, dd, $J$ = 8.0, 0.6 Hz), 7.71–7.69 (1H, m), 6.81 (2H, d, $J$ = 2.5 Hz), 6.74 (2H, d, $J$ = 8.7 Hz), 6.40 (2H, ddd, $J$ = 8.7, 2.5, 2.5 Hz), 5.49 (1H, ddd, $J$ = 9.2, 6.0, 3.4 Hz), 5.34 (1H, ddd, $J$ = 9.2, 6.0, 3.4 Hz), 5.06 (1H, d, $J$ = 3.8 Hz), 4.96 (1H, d, $J$ = 3.8 Hz), 4.25–4.16 (4H, m), 3.99–3.95 (3H, m), 3.92–3.89 (2H, m), 3.81–3.68 (5H, m), 3.63–3.55 (2H, m), 3.46 (1H, dd, $J$ = 9.7, 3.9 Hz), 3.27–3.22 (2H, m), 3.16 (1H, dd, $J$ = 9.8, 9.1 Hz), 0.65 (3H, s), 0.55 (3H, s).

$^{13}$C NMR (100 MHz, $d_4$-MeOD); $\delta_C$ 171.7, 169.2, 156.3, 151.9, 141.7, 138.1, 134.1, 129.8, 129.5, 128.99, 128.98, 126.7, 124.8, 117.48, 117.46, 114.37, 114.34, 95.3, 95.2, 93.7, 85.4, 83.4, 74.6, 74.3, 73.9, 73.6, 73.3, 73.2, 71.94, 71.92, 62.7, 60.7, 60.5, 42.3, 0.2, −1.4.

$^{19}$F{$^{1}$H} NMR (376.5 MHz, $d_4$-MeOD); $\delta_F$ -181.6.

HRMS (ESI); calc'd for C41H48F2N3O13Si [M+H+]+ : m/z 856.2919, found 856.2924.

**Synthesis of** $JF_{646}$**-Tre**

To a stirring solution of 6-trehalosamine (21mg, 62μmol, 1.5 equiv.)[43] in DMSO (2mL) was added $JF_{646}$ NHS ester (25mg, 42μmol, 1 equiv.).[37] The reaction was stirred at room temperature for 1 hour before being purified directly by prep-HPLC (10–90% MeCN/0.1% formic $acid_{(aq.)}$). Pure product fractions were then combined and lyophilized to give the title compound as a blue solid (22mg, 64%).

$^{1}$H NMR (400 MHz, $d_4$-MeOD); $\delta_H$ 8.04 (1H, dd, $J$ = 8.0, 1.3 Hz), 8.00 (1H, dd, $J$ = 8.0, 0.6 Hz), 7.71–7.69 (1H, m), 6.75 (2H, d, $J$ = 2.6 Hz), 6.70 (2H, d, $J$ = 8.7 Hz), 6.35 (2H, ddd, $J$ = 8.7, 2.5, 1.7 Hz), 5.06 (1H, d, $J$ = 3.8 Hz), 4.96 (1H, d, $J$ = 3.8 Hz), 3.97 (1H, ddd, $J$ = 10.0, 7.1, 2.9 Hz), 3.90 (8H, t, $J$ = 7.3 Hz), 3.81–3.68 (5H, m), 3.65–3.55 (2H, m), 3.46 (1H, dd, $J$ = 9.7, 3.8 Hz), 3.27–3.23 (2H, m), 3.16 (1H, dd, $J$ = 9.8, 9.1 Hz), 2.41–2.34 (4H, m), 0.63 (3H, s), 0.53 (3H, s).

$^{13}$C NMR (100 MHz, $d_4$-MeOD); $\delta_C$ 171.9, 169.3, 156.3, 152.89, 152.86, 141.6, 138.0, 133.39, 133.38, 130.0, 129 .4, 129.02, 129.00, 126.7, 124.8, 117.02, 116.99, 113.91, 113.87, 95.3, 95.2, 94.6, 74.6, 74.3, 73.9, 73.6, 73.3, 73.2, 72.0, 71.9, 62.7, 53.43, 53.41, 42.3, 17.8, 0.2, −1.4.

HRMS (ESI); calc'd for C41H50N3O13Si [M+H+]+ : m/z 820.3107, found 820.3105.

#### Figures

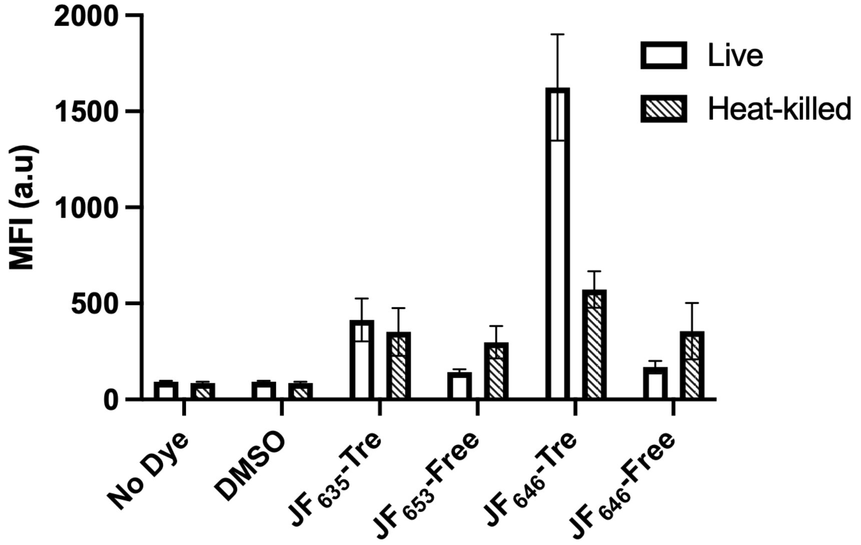


**Figure S1.** Flow cytometry raw MFI of live or heat-killed Msmeg cells incubated with 10 μM $JF_{635}$-Tre or 10 μM Free $JF_{635}$, 0.2% DMSO, or left untreated (No Dye) for 1 hour at 37˚C. Error bars denote standard deviation of three biological replicates. MFI (a.u.) = Mean Fluorescence Intensity (arbitrary units).

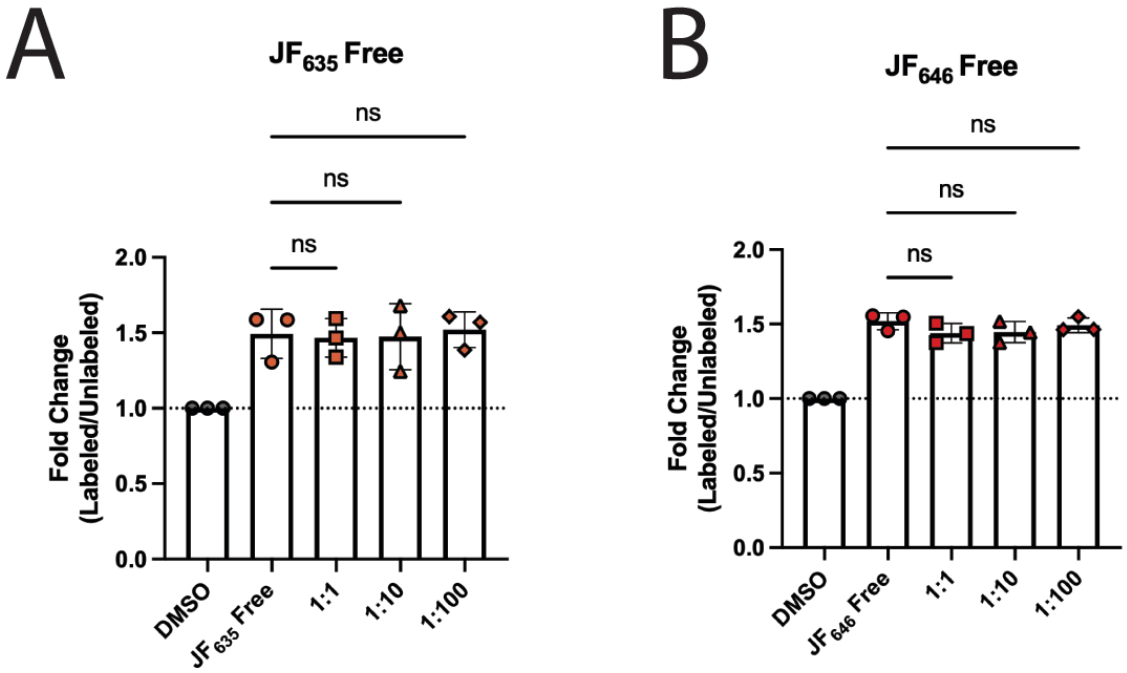


**Figure S2.** Flow cytometry MFI analysis of Msmeg cells incubated 1 hour with either (**A**) 10 μM unconjugated $JF_{635}$ ($JF_{635}$ Free), (**B**) 10 μM unconjugated $JF_{646}$ ($JF_{646}$ Free), or 0.2% DMSO for 1 hour at 37˚C with increasing amounts of exogenous trehalose. Error bars denote standard deviation of three biological replicates. $JF_{635}$ and $JF_{646}$ experiments were done in separate days. Error bars denote standard deviation of three biological replicates and data were analyzed by t-test or ANOVA tests in GraphPad Prism. MFI (a.u.) = Mean Fluorescence Intensity (arbitrary units).

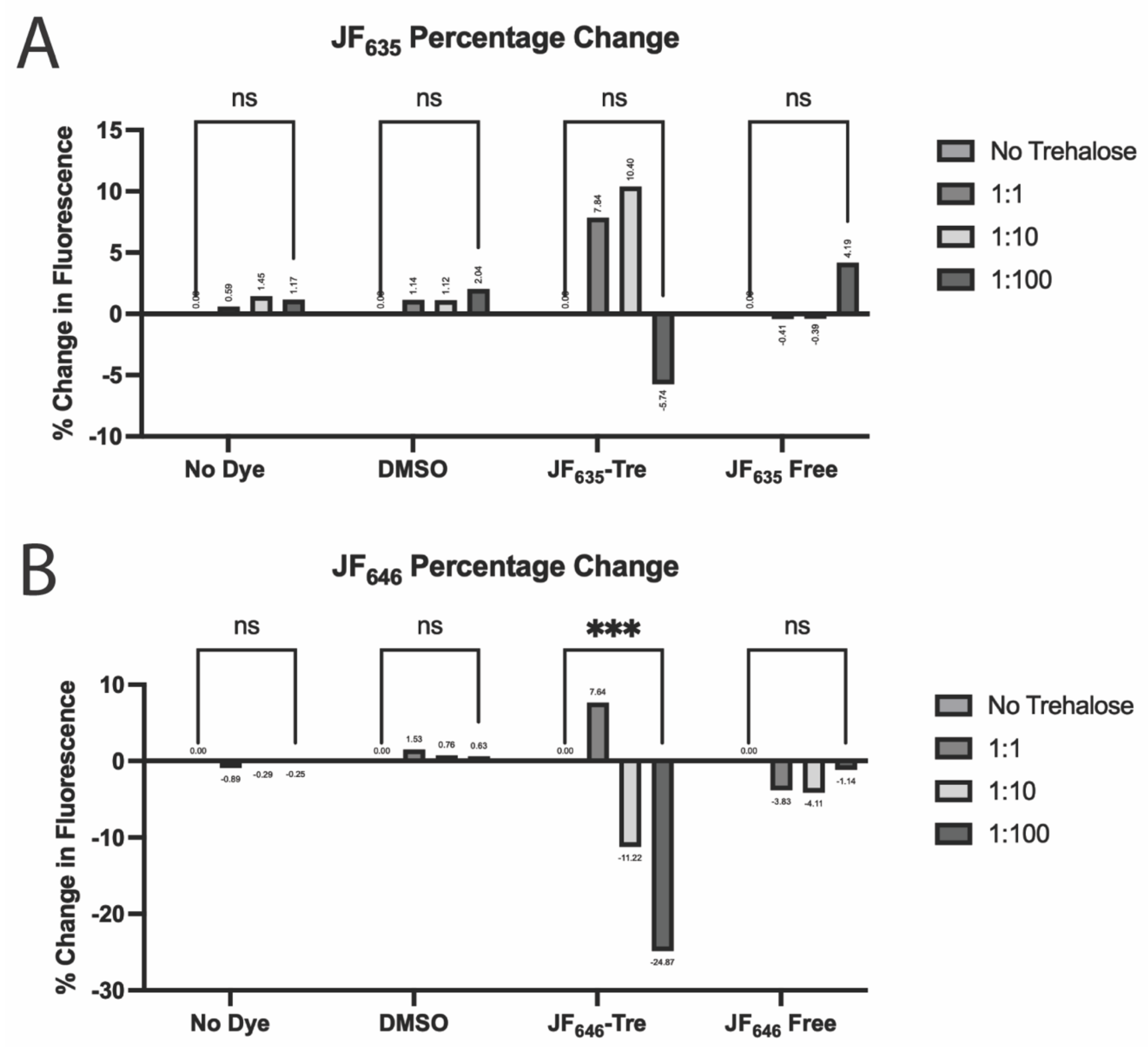


**Figure S3.** Percent change in fluorescence of Msmeg cells treated with (**A**) $JF_{635}$ or (**B**) $JF_{646}$. Percent change was calculated by taking the difference between the values of each condition and no exogenous trehalose condition (No Trehalose), dividing by the No Trehalose sample, and multiplying by 100. $JF_{635}$ and $JF_{646}$ experiments were done in separate days. Values reported are the mean of three biological replicates and data were analyzed by ANOVA tests in GraphPad Prism. MFI (a.u.) = Mean Fluorescence Intensity (arbitrary units). *p* values: *** < 0.001.

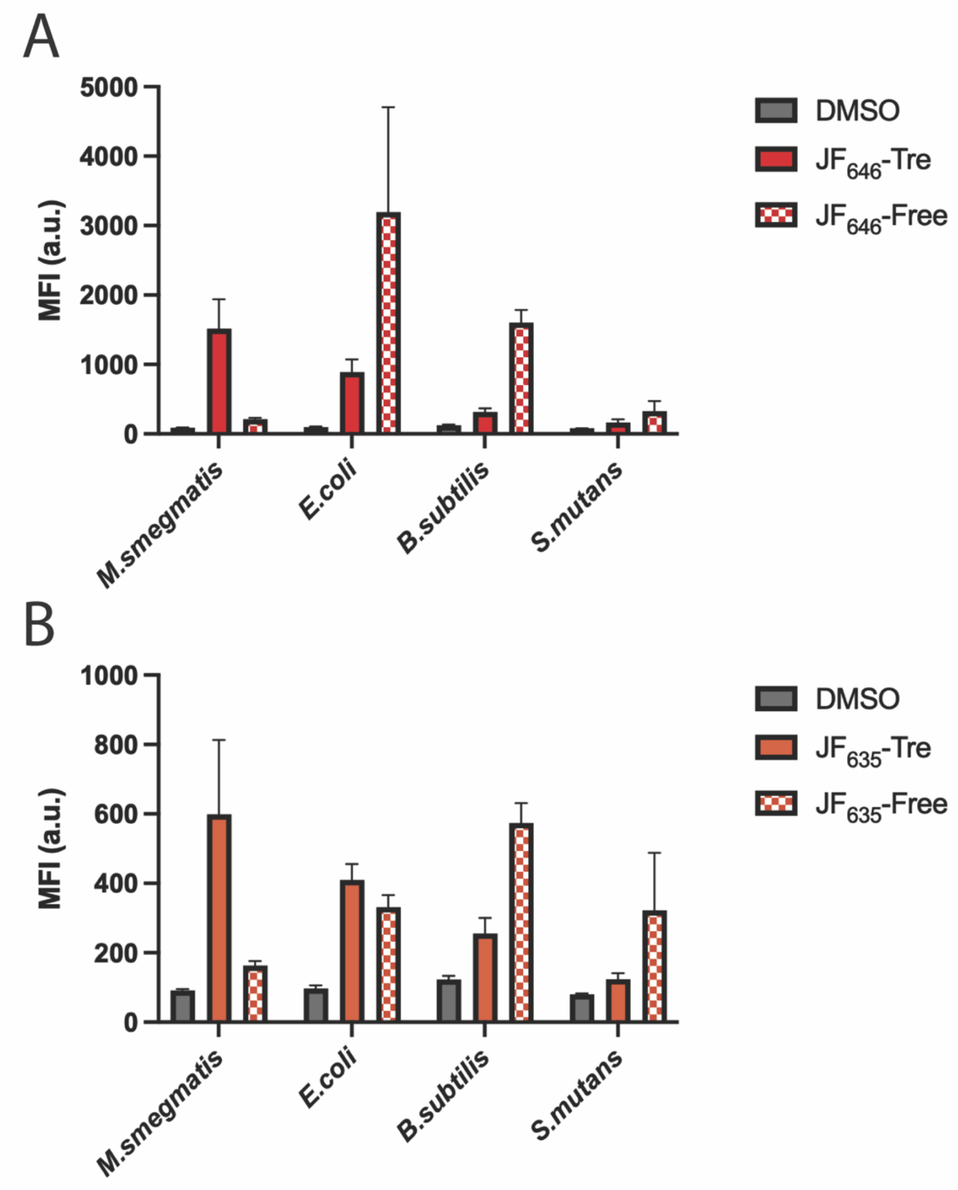


**Figure S4.** Flow cytometry raw MFI of Msmeg and non-mycobacterial species labeled with either (**A**) 10 µM $JF_{646}$-Tre, 10 µM unconjugated $JF_{646}$ ($JF_{646}$-Free), (**B**) 10 µM $JF_{635}$-Tre, 10 µM unconjugated $JF_{635}$ ($JF_{646}$-Free), or 0.2% DMSO. MFI (a.u.) = Mean Fluorescence Intensity (arbitrary units). Error bars denote standard deviation of three biological replicates.

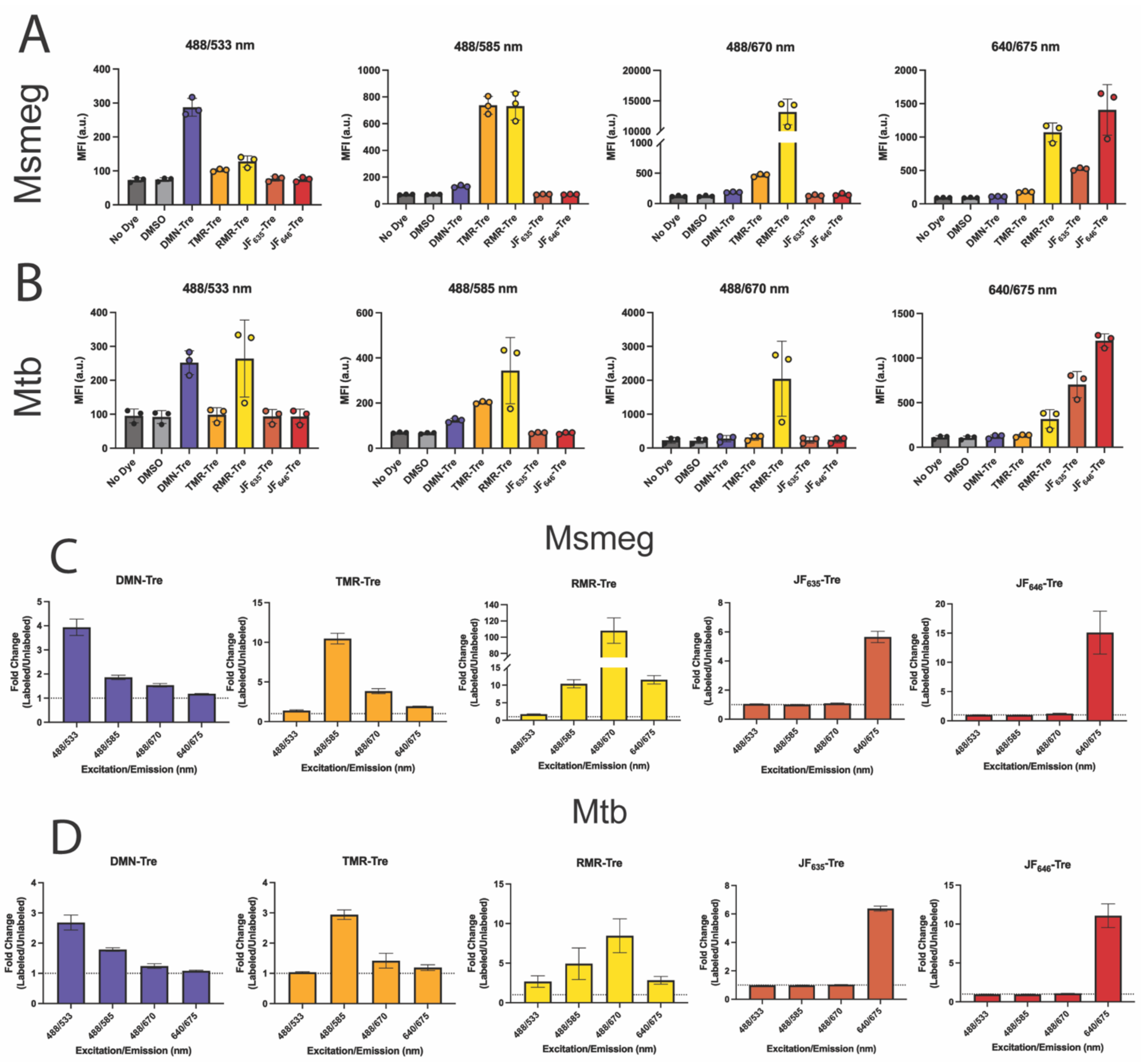


**Figure S5.** Raw fluorescence values of (**A**) Msmeg and (**B**) Mtb cultures by flow cytometry. Msmeg cultures incubated for 4 hours and Mtb cultures incubated for 16 hours at 37 ºC with 100 µM DMN-Tre, 100 µM TMR-Tre, 100 µM RMR-Tre, 10 µM $JF_{635}$-Tre, and 10 µM $JF_{646}$-Tre. (**C-D**) Fold-change analysis was performed afterwards, and results were arranged by probe with values for each excitation and emission pairs available in the instrument used. Fold change was calculated by taking the values of dye-labeled samples over no-dye control. Error bars denote standard deviation of three biological replicates.

### JF635-Tre ¹H NMR

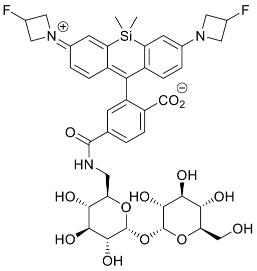


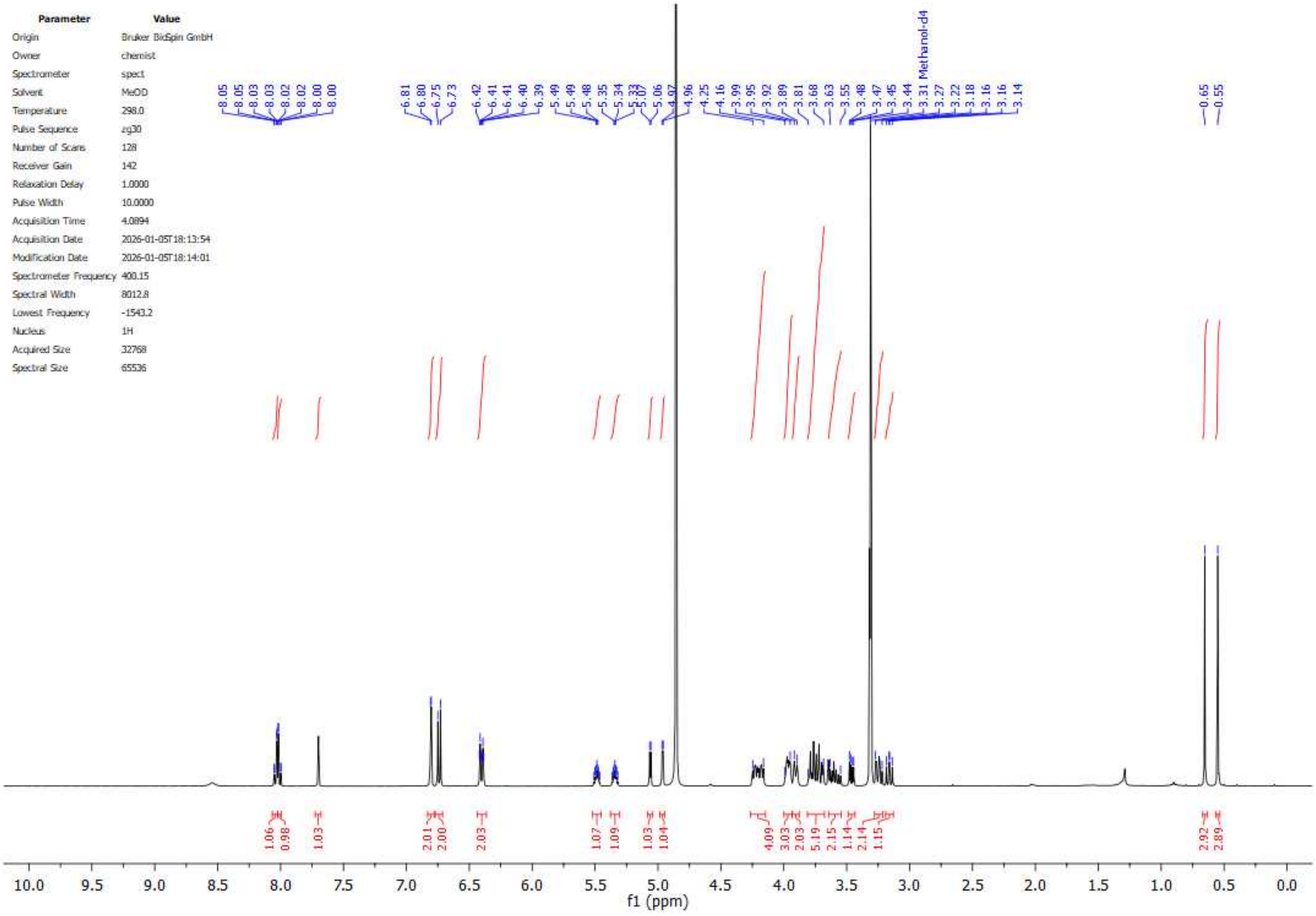

### JF635-Tre $^{13}C$ NMR

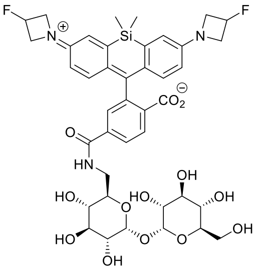


| Parameter | Value |
|---|---|
| Origin | Bruker BioSpin GmbH |
| Owner | chemist |
| Spectrometer | spect |
| Solvent | MeOD |
| Temperature | 298.0 |
| Pulse Sequence | zgpg30 |
| Number of Scans | 8192 |
| Receiver Gain | 197 |
| Relaxation Delay | 1.0000 |
| Pulse Width | 10.0000 |
| Acquisition Time | 2.3809 |
| Acquisition Date | 2026-01-06T02:08:56 |
| Modification Date | 2026-01-06T02:09:03 |
| Spectrometer Frequency | 100.62 |
| Spectral Width | 25000.0 |
| Lowest Frequency | -1292.2 |
| Nucleus | 13C |
| Acquired Size | 59522 |
| Spectral Size | 131072 |

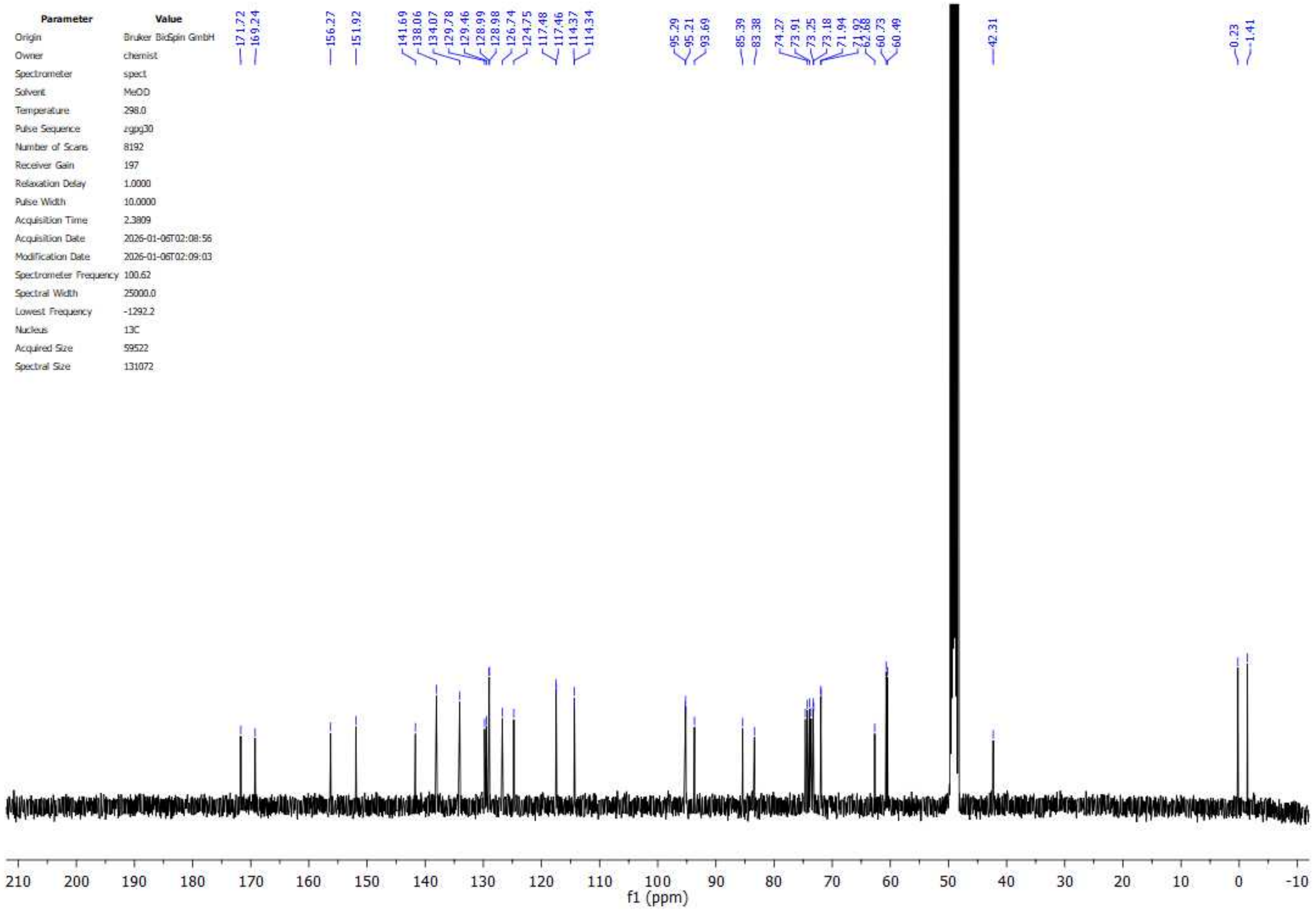

### JF635-Tre $^{19}$F NMR

| Parameter | Value |
|---|---|
| Origin | Bruker BioSpin GmbH |
| Owner | chemist |
| Spectrometer | spect |
| Solvent | MeOD |
| Temperature | 298.0 |
| Pulse Sequence | zgfhigqn.2 |
| Number of Scans | 64 |
| Receiver Gain | 197 |
| Relaxation Delay | 1.0000 |
| Pulse Width | 18.0000 |
| Acquisition Time | 0.7340 |
| Acquisition Date | 2026-01-05T18:17:19 |
| Modification Date | 2026-01-05T18:17:23 |
| Spectrometer Frequency | 376.52 |
| Spectral Width | 89285.7 |
| Lowest Frequency | -82294.6 |
| Nucleus | 19F |
| Acquired Size | 65536 |
| Spectral Size | 131072 |

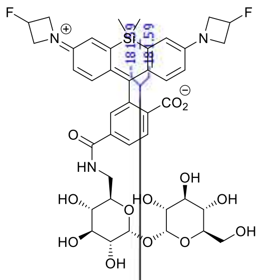


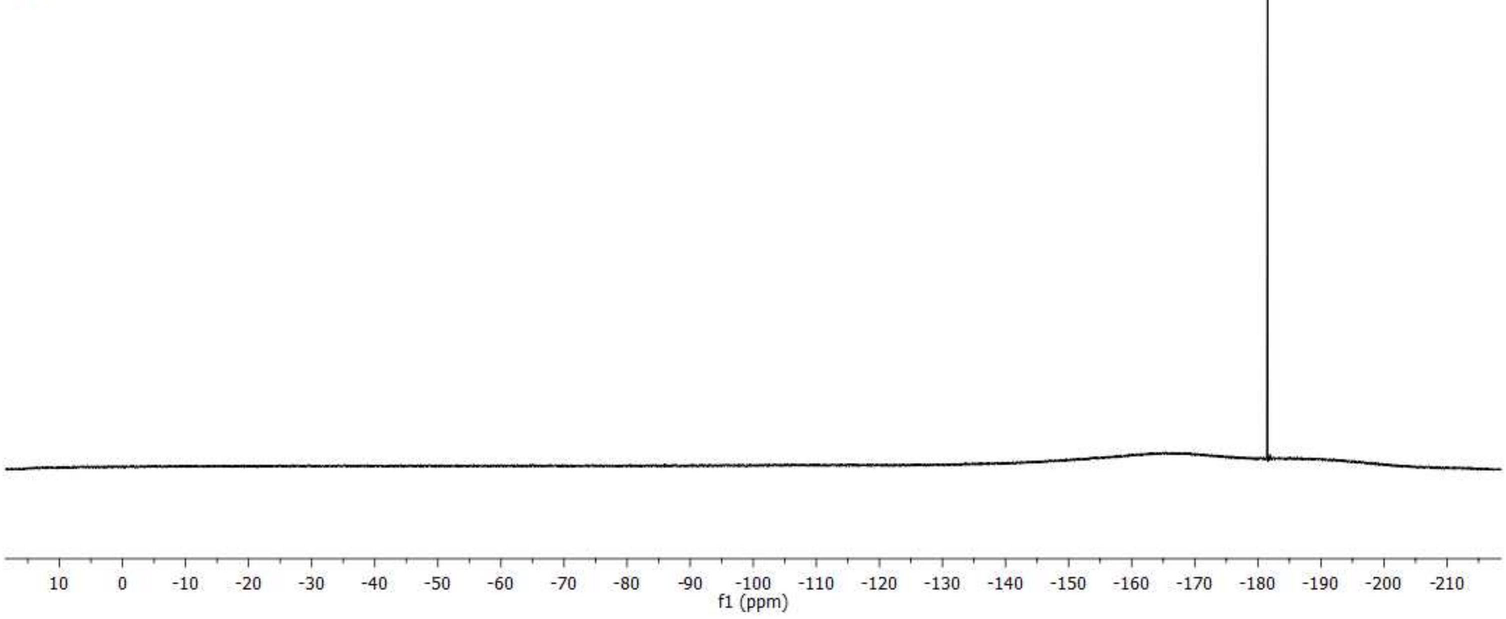

### JF646-Tre $^{1}$H NMR

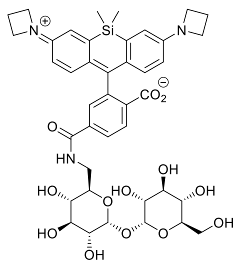


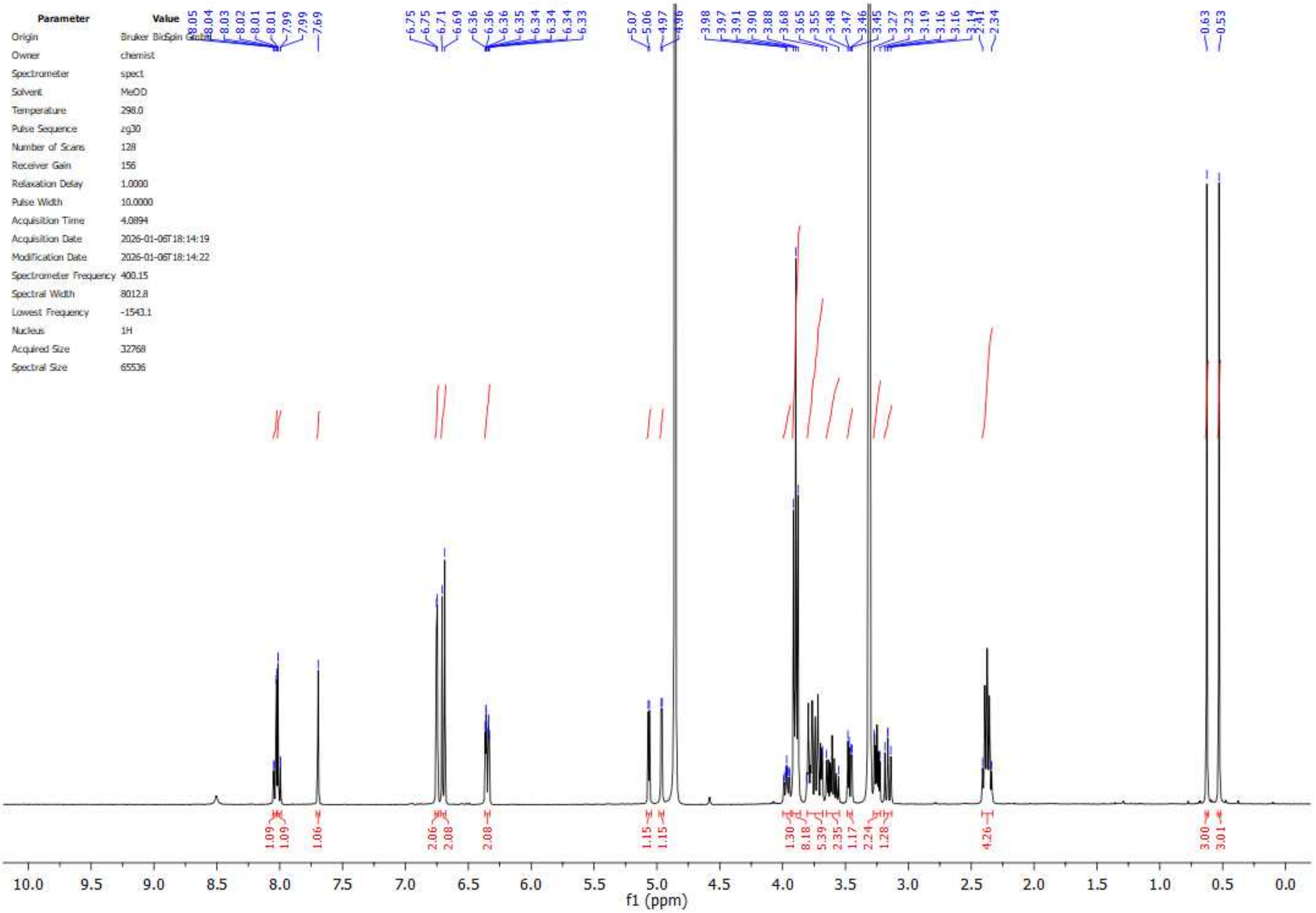


| Parameter | Value |
|---|---|
| Origin | Bruker BioSpin GmbH |
| Owner | chemist |
| Spectrometer | spect |
| Solvent | MeOD |
| Temperature | 298.0 |
| Pulse Sequence | zg30 |
| Number of Scans | 128 |
| Receiver Gain | 156 |
| Relaxation Delay | 1.0000 |
| Pulse Width | 10.0000 |
| Acquisition Time | 4.0894 |
| Acquisition Date | 2026-01-06T18:14:19 |
| Modification Date | 2026-01-06T18:14:22 |
| Spectrometer Frequency | 400.15 |
| Spectral Width | 8012.8 |
| Lowest Frequency | -1543.1 |
| Nucleus | 1H |
| Acquired Size | 32768 |
| Spectral Size | 65536 |

### JF646-Tre $^{13}C$ NMR

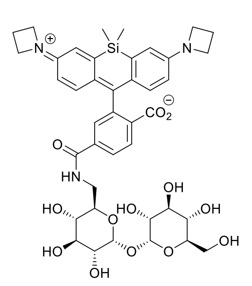


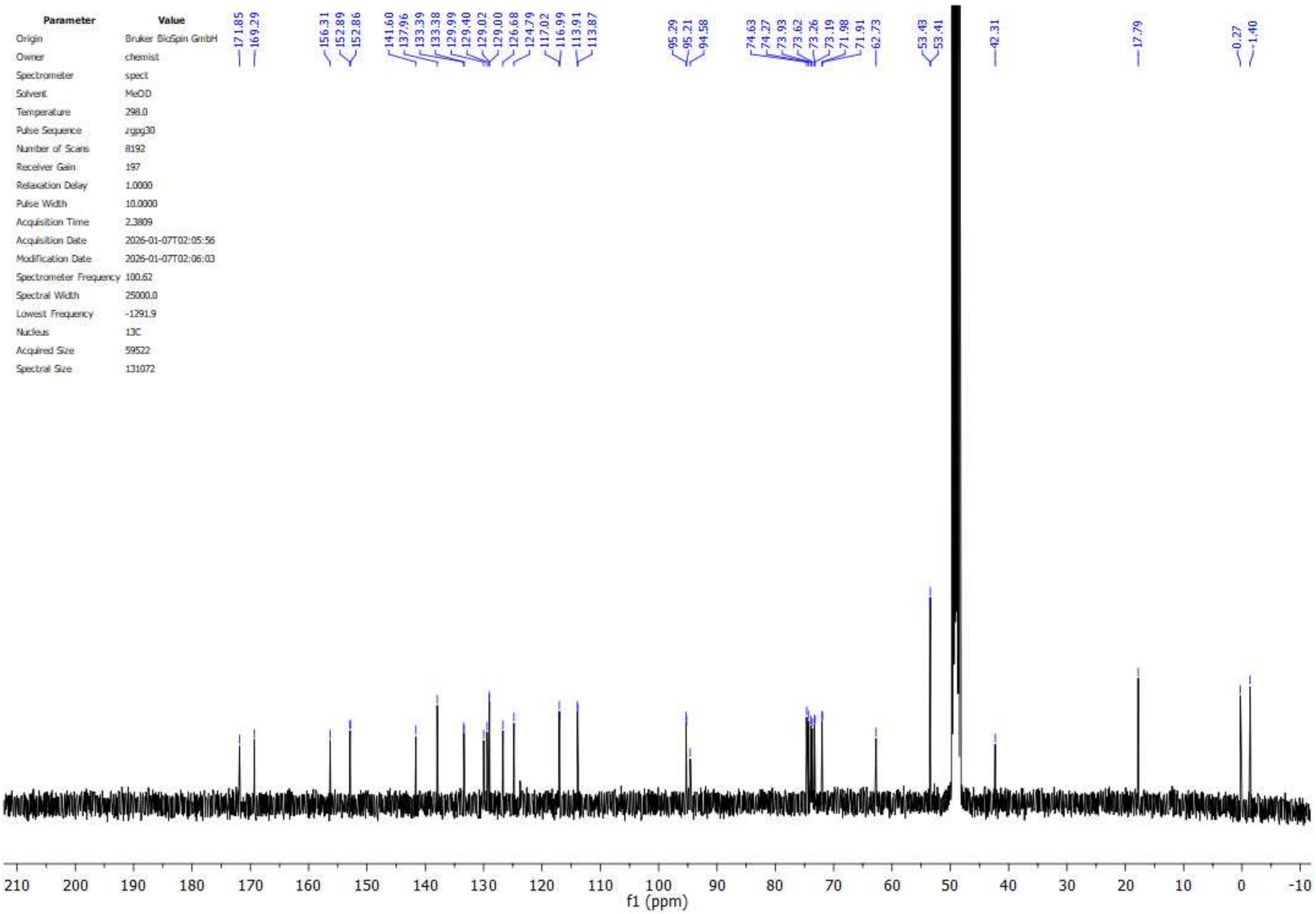
